# Plasma membrane protrusions mediate host cell-cell fusion induced by *Burkholderia thailandensis*

**DOI:** 10.1101/2022.02.24.481536

**Authors:** Nora Kostow, Matthew D. Welch

## Abstract

Cell-cell fusion is important for biological processes including fertilization, development, immunity, and microbial pathogenesis. Bacteria in the pseudomallei group of *Burkholderia* species, including *B. thailandensis*, spread between host cells by inducing cell-cell fusion. Previous work showed that *B. thailandensis*-induced cell-cell fusion requires intracellular bacterial motility and a bacterial protein secretion apparatus called the type VI secretion system-5 (T6SS-5), including the T6SS-5 protein VgrG5. However, the cellular level mechanism and T6SS-5 proteins important for bacteria-induced cell-cell fusion remained incompletely described. Using live cell imaging, we found bacteria used actin-based motility to push on the host cell plasma membrane to form plasma membrane protrusions that extended into neighboring cells. Then, membrane fusion occurred within these membrane protrusions, either proximal to the bacterium at the tip or elsewhere within a protrusion. Expression of VgrG5 by bacteria within membrane protrusions was required to promote cell-cell fusion. Furthermore, a second predicted T6SS-5 protein, TagD5, was also required for cell-cell fusion. In the absence of VgrG5 or TagD5, bacteria in plasma membrane protrusions were engulfed into neighboring cells. Our results suggest that the T6SS-5 effectors VgrG5 and TagD5 are secreted within membrane protrusions and act locally to promote membrane fusion.

## Introduction

Cell-cell fusion is important for biological processes including fertilization, development, and immunity (Chen et al., 2007). During cell-cell fusion, the plasma membranes of two cells are merged in a process that involves two key steps. In the first step, the two membranes, which are typically separated by extracellular components, are brought into close proximity (Chernomordik & Kozlov, 2003; Hernández & Podbilewicz, 2017). This often requires cellular factors such as the cytoskeleton and cell adhesion molecules (Zito et al., 2016; Hernández & Podbilewicz, 2017; Kim & Chen, 2019; Takito & Nakamura, 2020). In a second step, the remaining distance between the membranes is closed, the outer leaflets fuse to form a hemifusion intermediate, and the inner leaflets combine, resulting in the formation of a fusion pore without disrupting plasma membrane integrity (Chernomordik & Kozlov, 2003; Hernández & Podbilewicz, 2017). This requires the activity of proteins called fusogens (Chernomordik & Kozlov, 2003; Hernández & Podbilewicz, 2017). Once membrane fusion occurs, the small fusion pore then expands to generate one continuous cell (Hernández & Podbilewicz, 2017). Although the steps required for fusion are clear, the cellular and molecular level mechanisms are poorly understood. One approach to revealing cell-cell fusion mechanisms is to investigate microbe-induced cell-cell fusion processes.

The pseudomallei group of *Burkholderia* species are the only bacterial species known to directly induce cell-cell fusion (Kespichayawattana et al., 2000). This leads to the formation of multinucleated giant cells (MNGCs), both in cultured cells and in infected animals and human tissues (French et al., 2011; Harley et al., 1998; Kespichayawattana et al., 2000; West et al., 2008; K. T. Wong et al., 1995). Two species within this group, *B. pseudomallei* and *B. mallei*, cause the human disease melioidosis and equine disease glanders, respectively (Wiersinga et al., 2018; Wilkinson, 1981). A third species, *B. thailandensis*, is not thought to be a human pathogen and is used as a model system for studying aspects of infection with these pathogenic species (Haraga et al., 2008; West et al., 2008). *B. thailandensis* invades mammalian host cells, escapes the phagosome, and lives in the cytosol (Harley et al., 1998; Kespichayawattana et al., 2000). There, it undergoes intracellular bacterial actin-based (or flagellar) motility (French et al., 2011; Kespichayawattana et al., 2000; J. M. Stevens et al., 2005). Bacterial motility is important for efficient *B. thailandensis*-induced cell-cell fusion as a mutant deficient in both modes of motility induces cell-cell fusion with substantially reduced efficiency (French et al., 2011). Other bacterial pathogens, such as *Listeria monocytogenes* and *Rickettsia parkeri*, spread directly from cell-to-cell via a process that also involves actin-based motility (Lamason & Welch, 2017). Motility brings these bacteria to the plasma membrane where they enter into membrane protrusions that are engulfed into neighboring cells and are resolved into double membrane vesicles that they escape from to regain access to the cytosol (Tilney & Portnoy, 1989; Robbins et al., 1999; Monack & Theriot, 2001; Lamason et al., 2016; Lamason & Welch, 2017). *B. thailandensis* is also observed in plasma membrane protrusions (Kespichayawattana et al., 2000; J. M. Stevens et al., 2005; M. P. Stevens et al., 2005), but whether and how these protrusions contribute to cell-cell fusion is not known.

The second feature contributing to cell-cell fusion is a bacterial protein secretion apparatus called the type VI secretion system (T6SS) (Schell et al., 2007; Schwarz et al., 2010), a needle-like apparatus composed of a tube and a tip complex. To achieve secretion, bacterial proteins with effector functions can be translationally fused with T6SS needle tip components or can bind to T6SS tip or tube components (Jurėnas & Journet, 2021; Mougous et al., 2006; Pukatzki et al., 2006, 2007a). The T6SS secretes proteins by ejecting the tube and tip of the needle from the bacterium, a process that can puncture into a neighboring bacterium or host cell, releasing effector proteins into the target cell. Secretion can also occur without puncturing nearby cells by ejecting proteins into the extracellular environment (Jurėnas & Journet, 2021). *B. thailandensis* has five T6SS’s, of which T6SS-5 is the only T6SS necessary for pathogenesis in a mouse model of infection (Burtnick et al., 2011; Hopf et al., 2014; Pilatz et al., 2006; Schell et al., 2007; Schwarz et al., 2010). The T6SS-5 needle tip component VgrG5 is also required for cell-cell fusion (Schwarz et al., 2014; Toesca et al., 2014) and is the only protein known to be secreted by the T6SS-5 (Schwarz et al., 2014). VgrG5 contains a domain common to all VgrG proteins that trimerizes to form a blunt cone structure (Leiman et al., 2009; Spínola-Amilibia et al., 2016). Another component that is typically present at the T6SS tip is a PAAR (proline-alanine-alanine-arginine) family protein, which binds to the tip of a VgrG trimer resulting in an extended T6SS needle tip complex (Shneider, 2013). *B. thailandensis* encodes a PAAR protein within the T6SS-5 gene cluster called TagD5 (Lennings, West, et al., 2019). However, whether TagD5 is required for cell-cell fusion, and how the T6SS-5 tip components contribute to cell-cell fusion remain unknown.

To better understand the cellular pathway and bacterial factors leading to cell-cell fusion, we carried out live cell imaging of *B. thailandensis* as it induced cell-cell fusion. We found that cell-cell fusion occurred within host cell plasma membrane protrusions, with membrane fusion occurring both proximal to the bacterium at the protrusion tip or elsewhere in the protrusion. Expression of VgrG5 was required within membrane protrusions to promote cell-cell fusion. We also found that TagD5 was required for fusion. In the absence of VgrG5 or TagD5, bacterial protrusions were engulfed into neighboring cells. Our results suggest that the T6SS-5 effectors VgrG5 and TagD5 are secreted within membrane protrusions and act to promote cell-cell fusion.

## Results

### *B. thailandensis* induces cell-cell fusion at the tip or elsewhere within plasma membrane protrusions

To understand the cellular level mechanism by which *B. thailandensis* induces cell-cell fusion, we performed live cell imaging of cell-cell fusion events during *B. thailandensis* infection. We used a *B. thailandensis* strain deficient in flagellar motility (Δ*motA2*) but still competent for actin-based motility (hereafter called strain *Bt* WT) (French et al., 2011). For live cell imaging, we made a strain that also expressed GFP-tagged ClpV5 (ClpV5-GFP), a protein involved in disassembly of the T6SS in other bacteria (hereafter called strain *Bt*GFP WT) (Bonemann et al., 2009). ClpV5-GFP forms bright puncta in the bacterial cytosol of *B. thailandensis* (Lennings, Makhlouf, et al., 2019; Schwarz et al., 2014), allowing for clear visualization of the bacteria. Infections were carried out in monolayers of A549 human lung epithelial cells consisting of a 1:1 mixture of cells that stably expressed either an RFP plasma membrane marker (TagRFP-T-farnesyl) (Lamason et al., 2016) or stably expressed GFP in the cytosol. Upon infection and *B. thailandensis*-induced cell-cell fusion, MNGCs formed that expressed both the TagRFP-T-farnesyl plasma membrane marker and cytosolic GFP. For live cell imaging of cell-cell fusion, we observed bacteria that originated from an infected MNGC as they induced cell-cell fusion with a neighboring cell that expressed TagRFP-T-farnesyl plasma membrane marker but not cytosolic GFP. Therefore, as cell-cell fusion occurred, we observed the location of the bacterium relative to the RFP-labeled plasma membrane as well as the timing of cell-cell fusion as indicated by the diffusion of the cytosolic GFP from the MNGC into the cell that did not express GFP.

We observed that moving bacteria collided with the plasma membrane of the MNGC (termed the “donor” cell) and then moved into a membrane protrusion that extended into the neighboring cell (termed the “recipient” cell) (Figures 1 and 2, Videos S1 and S2). In the 144 cell-cell fusion events observed, we saw membrane protrusions in 141 events (in 3 events no plasma membrane protrusion was observed). Of these 141 events, 20 were selected for further analysis because the entire cell was visible, the process of protrusion formation and cell-cell fusion was captured from start to finish, and we could determine the location where cell-cell fusion was initiated. In 12/20 events, bacteria exited the protrusion at the protrusion tip and moved into the cytosol of the recipient cell (Figures 1 A and B, Videos S1-2). Shortly thereafter, the cytosolic GFP diffused from the donor cell into the recipient cell (Figures 1 A and B, Videos S1-2). This indicates that membrane fusion occurred at the tip of the membrane protrusion and formed a pore through which the bacteria moved (cartooned in Figure 1C). In the remaining 8/20 events, the bacteria remained at the plasma membrane protrusion tip, even as the GFP signal diffused from the donor cell into the recipient cell (Figures 2 A and B, Videos S3-4). In one example, based on the TagRFP-T-farnesyl signal, the protrusion clearly appeared to be separated from the donor MNGC yet still contained the bacterium (Figure 2D, Video S3). Therefore, in these examples, the membrane fusion occurred at a distance from the bacteria (cartooned in Figure 2C). In many events (n = 14), after the GFP diffused from the donor cell into the recipient cell, the now merged plasma membrane spread apart in the area where the protrusion had formed (Figures 1 and 2, Videos S1-4, Supplemental Figure 1). This indicates that membrane fusion occurred within the membrane protrusion and expanded, leading to a continuous cytoplasm between the donor and recipient cell and expanding the size of the MNGC. These observations indicate that cell-cell fusion occurs within membrane protrusions, either at the protrusion tip or elsewhere in the protrusion.

**Figure 1:**
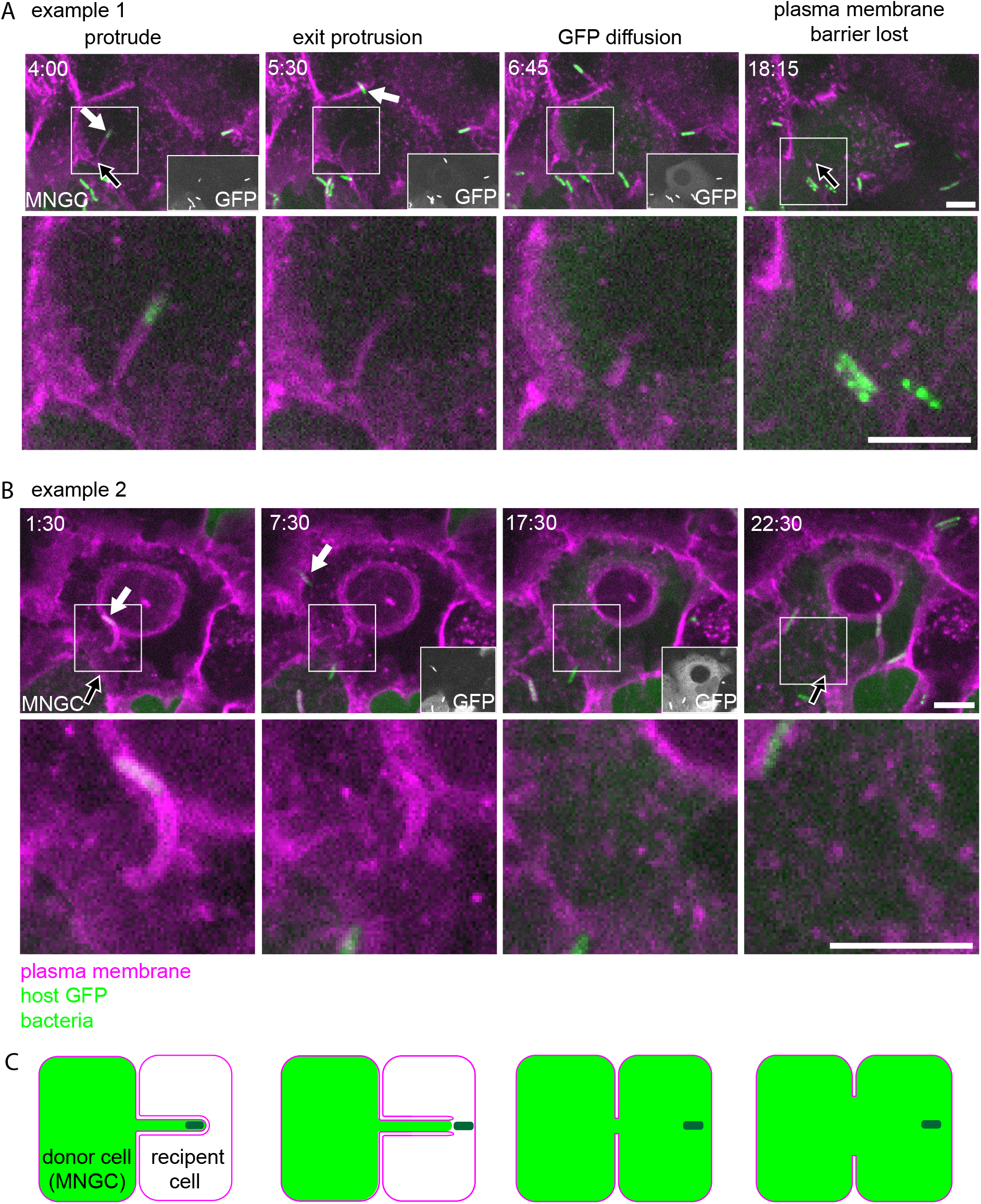
In one observed pathway, *B. thailandensis* spreads by inducing cell-cell fusion at the protrusion tip. (A and B) Live-cell imaging stills of two examples of *Bt*GFP WT while inducing cell-cell fusion. A 1:1 mixture of A549 cells that expressed the plasma membrane marker TagRFP-T-farnesyl or cytoplasmic GFP were used. Times represent min:s post protrusion formation. Images taken at ~16 h post infection. Scale bars are 5 μm. White arrows highlight the bacterium forming the protrusion. Black arrows highlight the region of protrusion entry. Where GFP signal is difficult to see, insets with increased brightness are shown. (C) Model of cell-cell fusion occurring at the protrusion tip.

**Figure 2:**
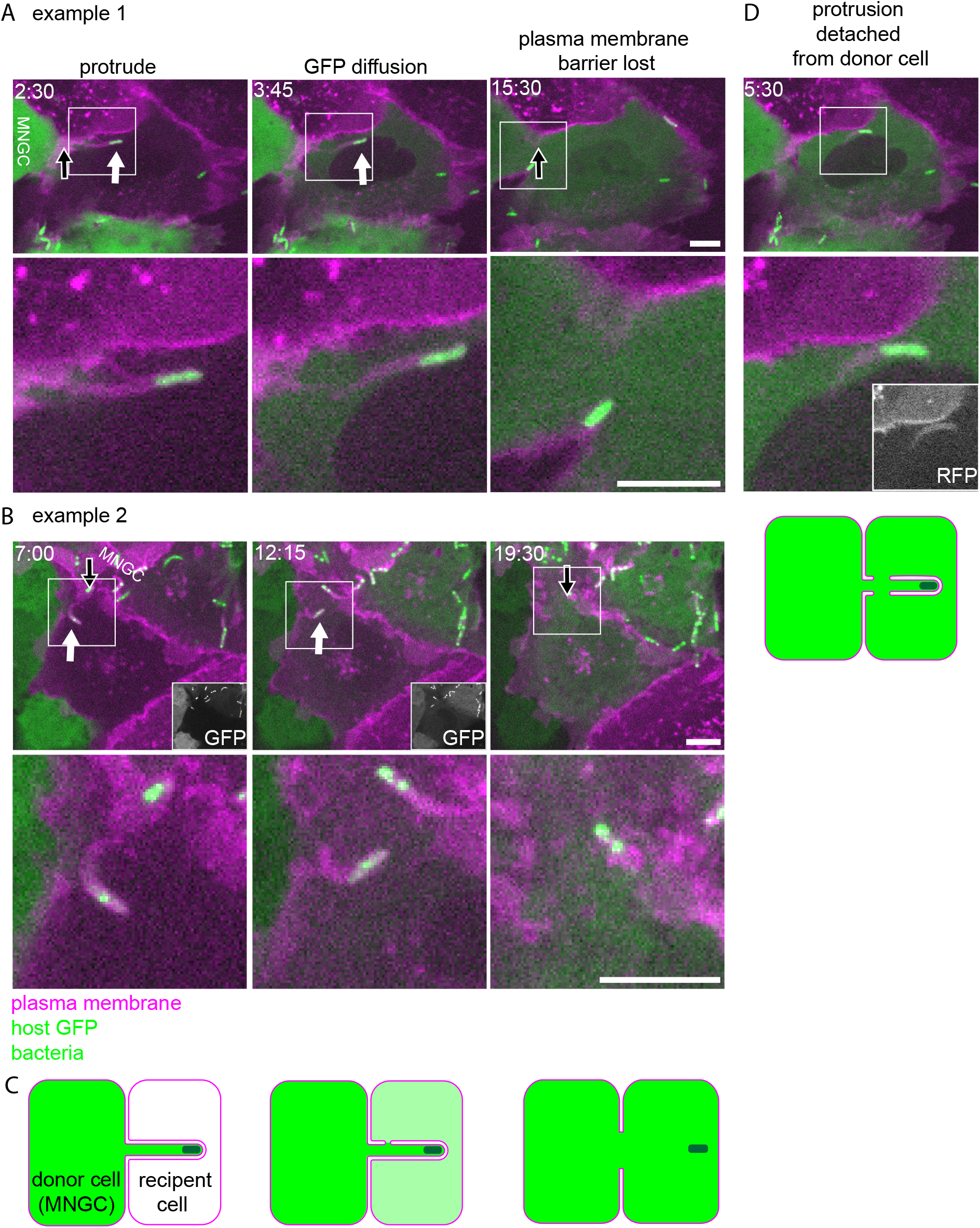
In one observed pathway, *B. thailandensis* spreads by inducing cell-cell fusion elsewhere within the protrusion. (A and B) Live-cell imaging stills of two examples of *Bt*GFP WT while inducing cell-cell fusion. A 1:1 mixture of A549 cells that expressed the plasma membrane marker TagRFP-T-farnesyl or cytoplasmic GFP were used. Times represent min:s post protrusion formation. Images taken at ~16 h post infection. Scale bars are 5 μm. White arrows highlight the bacterium forming the protrusion. Black arrows highlight the region of protrusion entry. Where GFP signal is difficult to see, insets with increased brightness are shown. (C) Model of cell-cell fusion occurring elsewhere within the protrusion. (D) Still showing visible detachment of the bacterium-containing protrusion from the donor cell.

To understand how protrusion morphology and timing might contribute to the cell-cell fusion pathway, we measured the maximum protrusion length and timing of cell-cell fusion events. The length of protrusions at their maximum was 8 +/− 2 μm (Figure 3A) (all experimentally determined values from this study are listed as mean +/− SD), shorter than the length previously observed for protrusions induced by *L. monocytogenes* (~17 μm on average) but longer than those induced by *R. parkeri* (~3 μm on average) (Lamason et al., 2016). There was no difference in maximum protrusion length during cell-cell fusion events that occurred at the protrusion tip versus elsewhere in the protrusion (Figure 3B), indicating that differences in the location of fusion are not related to protrusion length. To determine how long it takes for *B. thailandensis* to induce cell-cell fusion, we quantified the time from the start of protrusion formation to the time at which cytosolic GFP diffused into the recipient cell. This time was 8 +/− 3 min (Figure 3C; these data likely overrepresent shorter events due to the experimental difficulty of capturing long events) and there was no correlation between maximum protrusion length and the speed at which cell-cell fusion occurred (Figure 3D, R^2^=0.09). When binned based on the location of membrane fusion, membrane fusion at the protrusion tip occurred slightly faster than membrane fusion that occurred elsewhere in the protrusion (Figure 3E). *B. thailandensis*-induced cell-cell fusion occurred with similar or faster timing compared with the protrusion uptake pathway of *R. parkeri* (~10 minutes) and *L. monocytogenes* (~20 minutes) (Lamason et al., 2016). Therefore, *B. thailandensis*-induced cell-cell fusion appears to occur quickly compared with the cell-to-cell spread processes of other bacteria.

**Figure 3:**
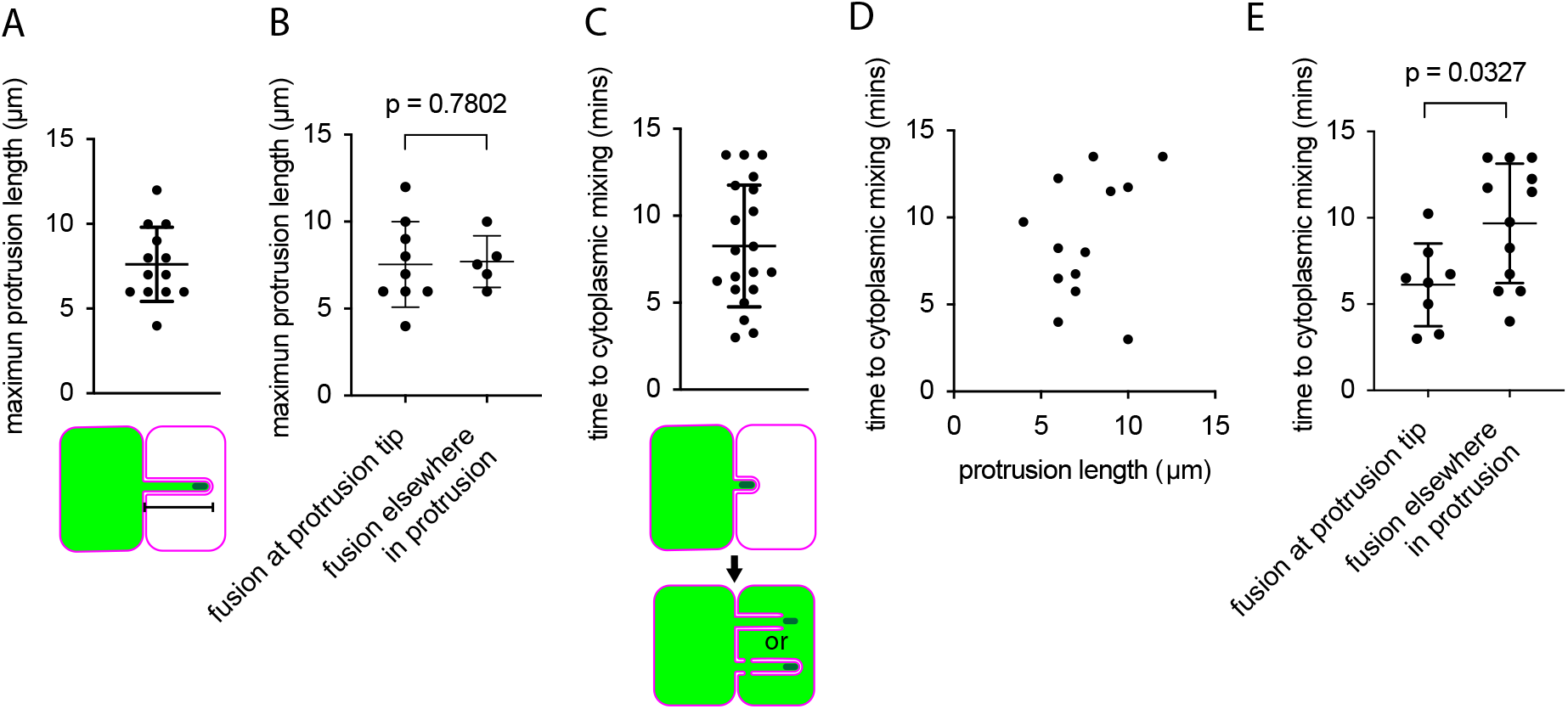
Quantification of *B. thailandensis* inducing cell-cell fusion live cell imaging dataset. (A) Graph of maximum protrusion length from videos where entire protrusion was visible (n=12). (B) Graph of maximum protrusion length for membrane fusion that occurred at the protrusion tip (n=9) versus elsewhere in the protrusion (n=4). (C) Graph of time to cytoplasmic mixing (n=20). (D) Graph of time to cytoplasmic mixing versus maximum protrusion length (n=12). R^2^ = 0.09197, p = 0.3138. (E) Graph of time to cytoplasmic mixing for membrane fusion that occurred at the protrusion tip (n=8) versus elsewhere in the protrusion (n=12). For (A-D,E) P values were calculated by unpaired Mann-Whitney test, data are mean +/− SD.

### VgrG5 acts at the membrane fusion step

The T6SS tip protein VgrG5 was previously found to be necessary for *B. thailandensis*-induced cell-cell fusion (Schwarz et al., 2014; Toesca et al., 2014). However, it was unknown at which stage of the cell-cell fusion pathway VgrG5 contributes. To investigate this, we generated identical a *ΔvgrG5* deletion mutants in both *Bt* (*Bt* Δ*vgrG5*) and *Bt*GFP (*Bt*GFP Δ*vgrG5*) strain backgrounds. We confirmed that *Bt*GFP Δ*vgrG5* did not express VgrG5 by western blotting using an anti-VgrG5 antibody we generated (Supplemental Figure 2). We then infected A549 cells that expressed TagRFP-T-farnesyl with a Δ*vgrG5* deletion mutant in the *Bt*GFP strain (*Bt*GFP Δ*vgrG5*) and performed live cell imaging. We found that *Bt*GFP Δ*vgrG5* formed membrane protrusions (Figure 4A, Video S5) that appeared similar to protrusions formed by *Bt*GFP WT (Figures 1 and 2). Rather than inducing cell-cell fusion, *Bt*GFP Δ*vgrG5* bacteria in protrusions were instead engulfed into the recipient cell (Figure 4A, Video S5). Because this engulfment pathway is similar to the process that occurs during *R. parkeri* and *L. monocytogenese* cell-to-cell spread (Lamason & Welch, 2017), protrusions formed by *Bt*GFP Δ*vgrG5* are likely engulfed into double membrane vacuoles (cartooned in Figure 4B). These bacteria remained in these vacuoles for the duration of the imaging session. However, even though we did not observe such events, some bacteria still accessed the host cell cytosol after engulfment into recipient cells because some bacteria underwent actin-based motility as evidenced by their presence in plasma membrane protrusions of secondary cells (Supplemental Figure 3). These results indicate that VgrG5 is specifically involved in the membrane fusion step of the cell-cell fusion pathway.

**Figure 4:**
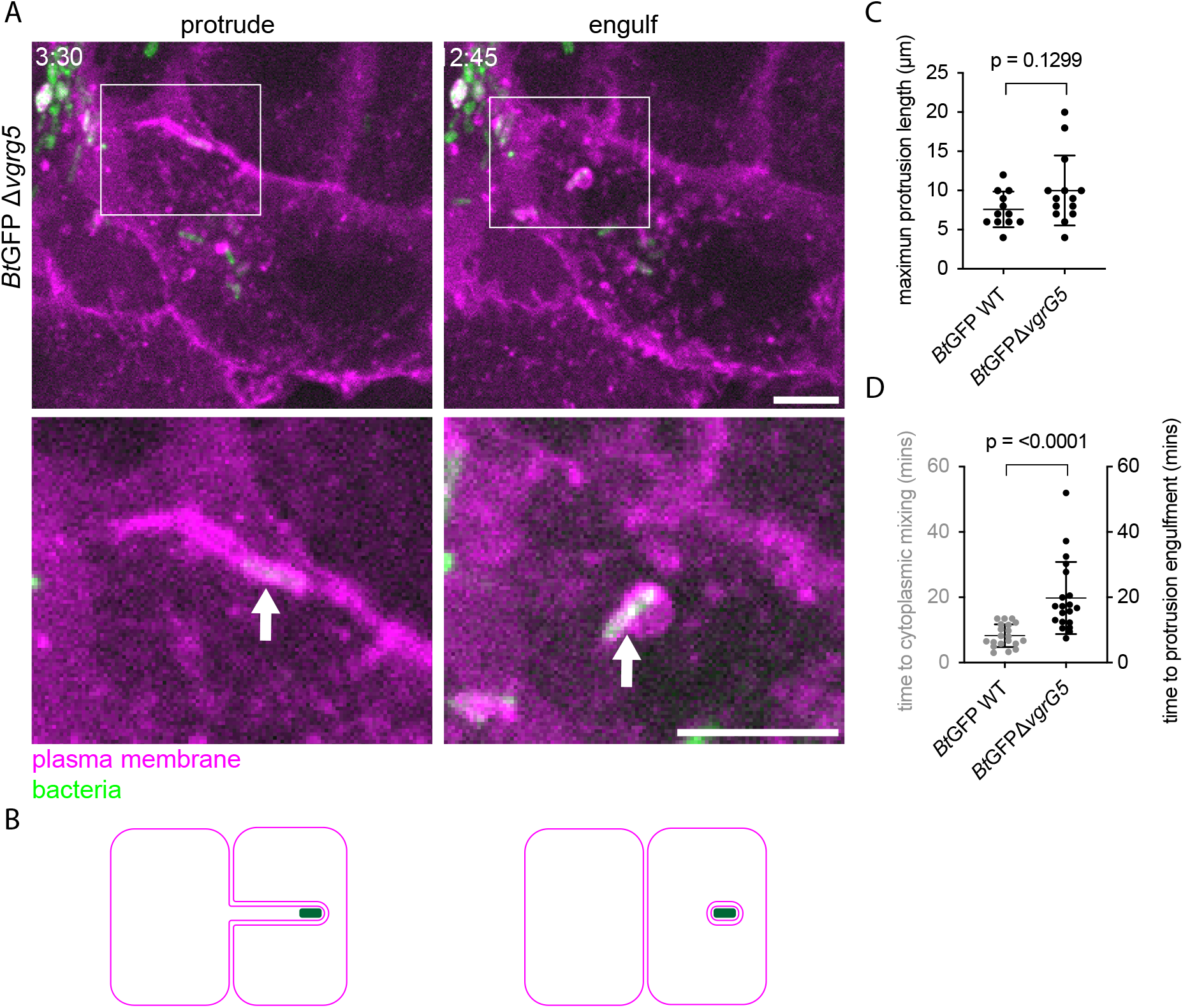
VgrG5 acts at the membrane fusion step. (A) Live cell imaging stills of *Bt*GFP Δ*vgrG5* during cell-to-cell spread. A549 cells that expressed TagRFP-T-farnesyl were used. Times represent min:s post protrusion formation. All images taken at ~24 h post infection. Scale bars are 5 μm. (B) Model of spread. (C) Graph of maximum protrusion length for *Bt*GFP WT (n=12) and *Bt*GFP Δ*vgrG5* (n=14). (D) Graph of time to cytoplasmic mixing (n=20) or protrusion engulfment (n=20). For (C-D), P values were calculated by unpaired Mann-Whitney tests, data are mean +/− SD.

To further compare the non-canonical cell-to-cell spread of *Bt*GFP Δ*vgrG5* with *Bt*GFP WT-induced cell-cell fusion, we measured the maximum protrusion lengths and timing of cellcell fusion or engulfment for both strains. Maximum protrusion lengths were not significantly different between *Bt*GFP WT and *Bt*GFP Δ*vgrG5* (Figure 4C). This suggests that VgrG5 does not contribute to protrusion formation. Compared with the time it took for *Bt*GFP WT to induce cell-cell fusion, the engulfment of *Bt*GFP Δ*vgrG5* took significantly longer (Figure 4D). *Bt*GFP WT in membrane protrusions were also occasionally engulfed (Supplemental Figure 4, Video S6). *Bt*GFP WT engulfment took significantly longer than *Bt*GFP WT-induced cell-cell fusion (Supplemental Figure 4B) and was not significantly different than engulfment of *B*tGFP Δ*vgrG5* (Supplemental Figure 4C). Our finding that *B*tGFP Δ*vgrG5* is engulfed into recipient cells, and that engulfment occurs more slowly than cell-cell fusion, suggests that inducing cell-cell fusion overrides a slower default engulfment pathway.

### *B. thailandensis* must express VgrG5 within a protrusion to induce cell-cell fusion

Having determined that membrane fusion can occur at a distance from the bacterium and that VgrG5 functions at the membrane fusion step of the cell-cell fusion pathway, we wondered whether VgrG5 could be supplied by other bacteria elsewhere in an infected cell. To answer this question, we performed a co-infection experiment in monolayers of A549 cells that consisted of a mixture of cells that stably expressed either TagRFP-T-farnesyl or stably expressed GFP in the cytosol. We co-infected these monolayers with *Bt* WT that expressed BFP (*Bt*BFP WT) (Benanti et al., 2015) and *Bt*GFP Δ*vgrG5*. We then performed live cell imaging, with a focus on *Bt*GFP Δ*vgrG5* bacteria that formed protrusions from MNGC donor cells that extended into neighboring recipient cells that expressed TagRFP-T-farnesyl (Figure 5A). If VgrG5 supplied by *Bt*BFP WT could rescue the ability of *Bt*GFP Δ*vgrG5* to induce cell-cell fusion, then we would observe diffusion of the GFP signal due to cell-cell fusion (Figure 5A, top). Alternatively, if VgrG5 supplied by *Bt*BFP WT could not rescue the ability of *Bt*GFP Δ*vgrG5* to induce cell-cell fusion, *Bt*GFP Δ*vgrG5* in membrane protrusions would be engulfed (Figure 5A, bottom). In all 10 instances observed, *Bt*GFP Δ*vgrG5* membrane protrusions were engulfed by the recipient cell and no cytosolic GFP diffused into the recipient cell during engulfment (Figure 5B, Video S7). This observation suggests that VgrG5 must be expressed by bacteria within membrane protrusions to promote cell-cell fusion.

**Figure 5:**
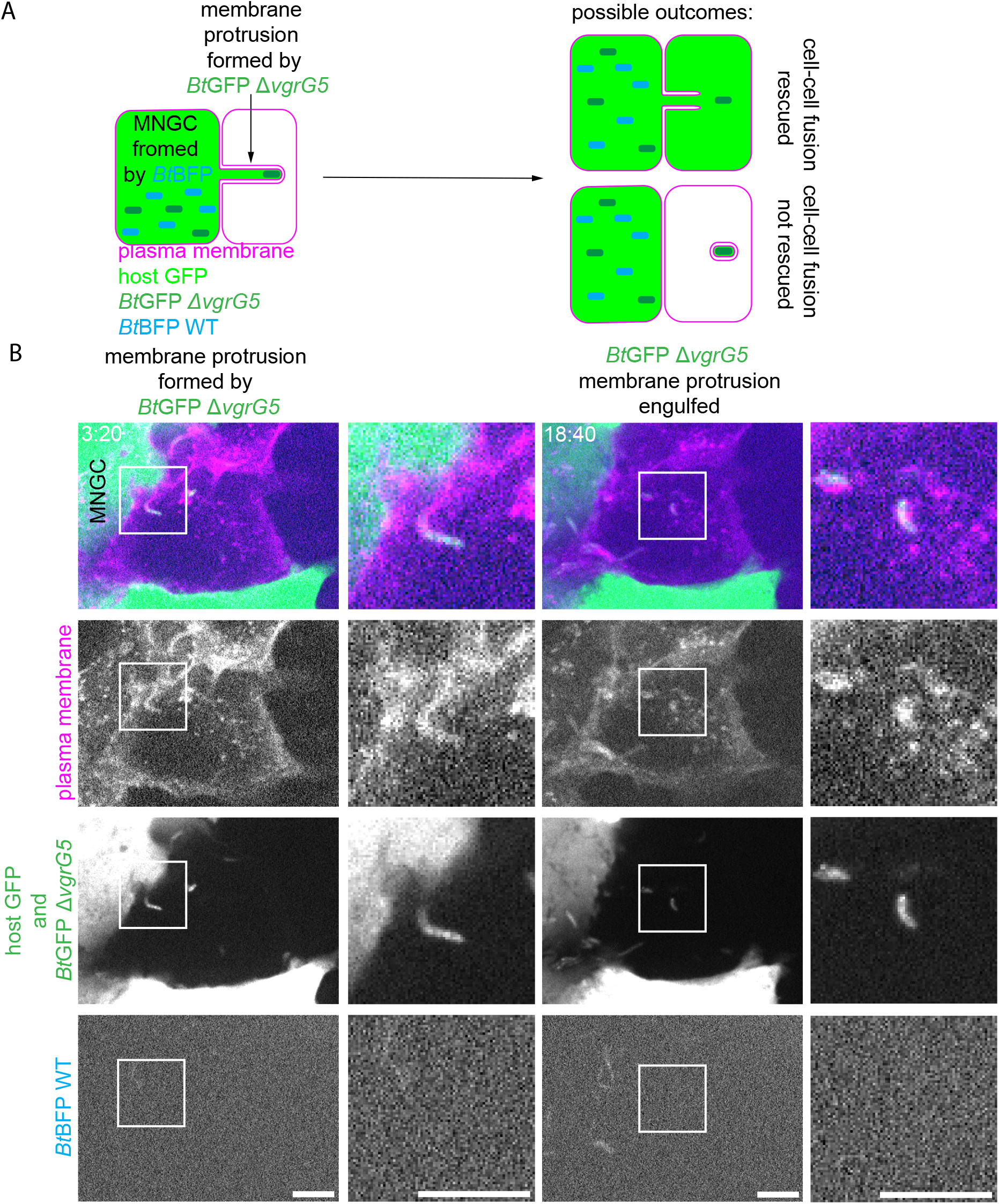
*B. thailandensis* must secrete VgrG5 within a protrusion to induce cell-cell fusion. (A) Experimental design and possible outcomes. (B) Live cell imaging stills of *Bt*GFP Δ*vgrg5* spreading from an MNGC initially formed by cell-cell fusion induced by *Bt*BFP WT bacteria. Times represent min:s after the video began. Scale bars are 5 μm.

### TagD5 is required for inducing cell-cell fusion and acts at the membrane fusion step

The PAAR protein TagD5 is encoded in the same T6SS-5 gene cluster as VgrG5 (Burtnick et al., 2011; Hopf et al., 2014; Lennings, West, et al., 2019; Pilatz et al., 2006; Schwarz et al., 2010), and based on the known interaction between VgrG and PAAR proteins (Shneider, 2013), we hypothesized that it functions with VgrG5 to induce membrane fusion. To test this, we generated identical Δ*tagD5* deletion mutants in both *Bt* (*Bt* Δ*tagD5*) and *Bt*GFP (*Bt*GFP Δ*tagD5*) strain backgrounds. We investigated whether *Bt*GFP Δ*tagD5* expressed VgrG5 by western blotting using our anti-VgrG5 antibody and found that *Bt* Δ*tagD5* exhibited reduced levels of VgrG5 protein (Supplemental Figure 2). Therefore, TagD5 influences VgrG5 expression or stability.

To test whether TagD5 is required for cell-cell fusion, we first employed a plaque size assay, which was previously used to determine the extent of cell-cell fusion (Benanti et al., 2015; French et al., 2011), in Vero cells. *Bt* Δ*tagD5* failed to form a plaque, as did *Bt* Δ*vgrG5* (Figure 6A). This is consistent with functions for both VgrG5 and TagD5 in cell-cell fusion.

**Figure 6:**
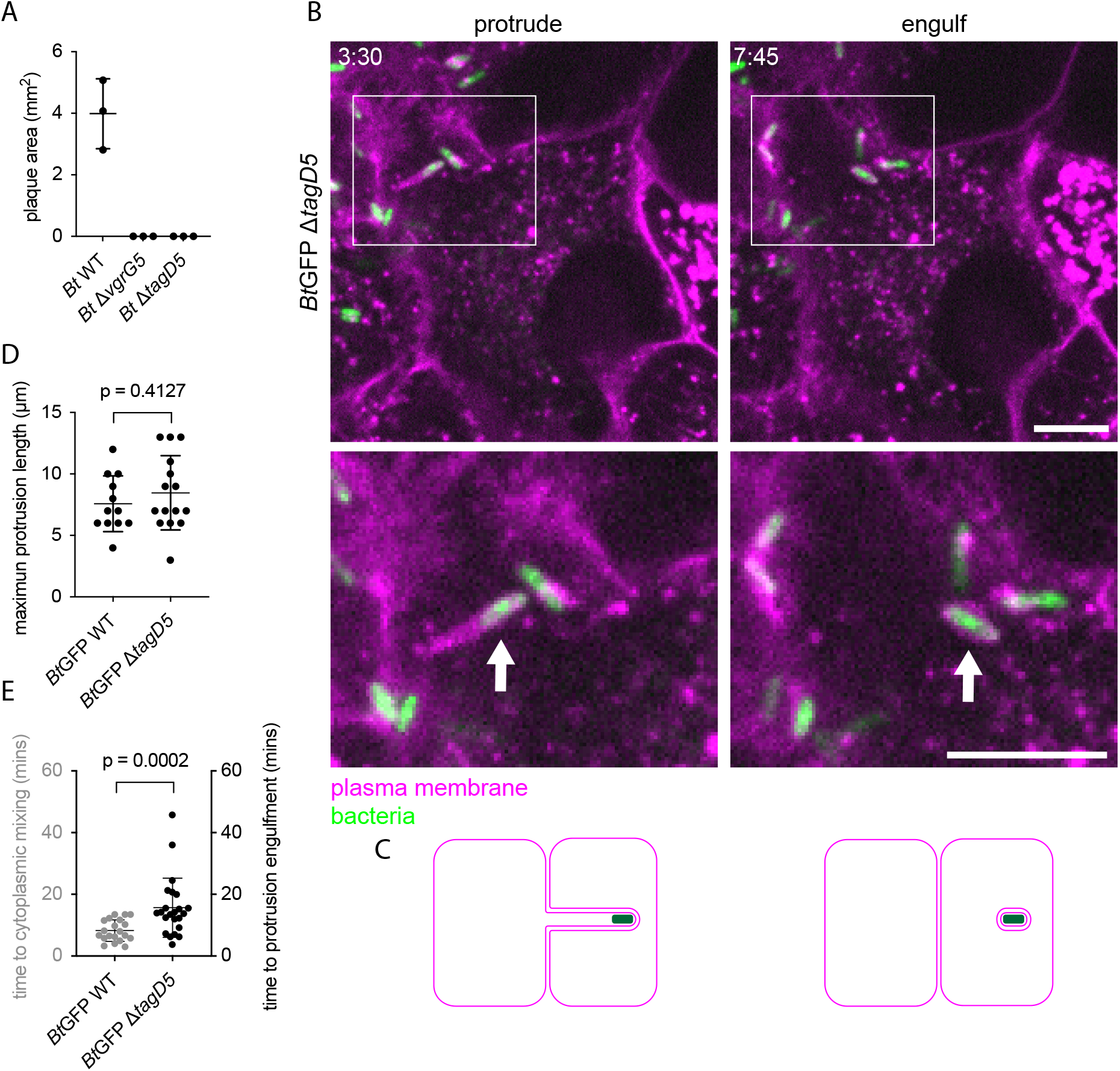
TagD5 is required for inducing cell-cell fusion and acts at the membrane fusion step. (A) Plaque areas of Vero cells infected with the indicated strains. N=3 experiments, 9-11 plaques per experiment. (B) Live-cell imaging stills of *Bt*GFP Δ*tagD5* during cell-to-cell spread. A549 cells that expressed TagRFP-T-farnesyl were used. Times represent min:s post protrusion formation. All images taken at ~24 h post infection. Scale bars are 5 μm. (C) Model of spread. (D) Graph of maximum protrusion length for *Bt*GFP WT (n=12) and *Bt*GFP Δ*tagD5* (n=15). (E) Graph of time to cytoplasmic mixing (n=20) or protrusion engulfment (n=23). For (A, D-E) P values were calculated by unpaired Mann-Whitney tests, data are mean +/− SD.

To determine the step at which TagD5 acts in the cell-cell fusion pathway, we next performed live cell imaging of A549 cells that expressed TagRFP-T-farnesyl infected with *Bt* Δ*tagD5* made in the *Bt*GFP strain (*Bt*GFP Δ*tagD5*). The phenotypes exhibited by *Bt*GFP Δ*tagD5* were nearly identical to those of *Bt*GFP Δ*vgrG5*. *Bt*GFP Δ*tagD5* did not induce cell-cell fusion and instead formed membrane protrusions that were engulfed by the recipient cell (Figure 6B,C, Video S8). Maximum protrusion lengths were not significantly different between *Bt*GFP WT and *Bt*GFP Δ*tagD5* (Figure 6D), similar to *Bt*GFP Δ*vgrG5* (Figure 4C). Furthermore, compared with the time it took for *Bt*GFP WT to induce cell-cell fusion, the engulfment of *Bt*GFP Δ*tagD5* took significantly longer (Figure 6E), similar to engulfment of *Bt*GFP Δ*vgrG5* (Figure 4D). There was no difference between the time to engulfment of *Bt*GFP Δ*vgrG5* and *Bt*GFP Δ*tagD5* (Supplemental Figure 4C). Therefore, both TagD5 and VgrG5 are required for the membrane fusion step of the cell-cell fusion pathway, consistent with them working together during this step.

## Discussion

Here we describe the cellular pathway leading to host cell-cell fusion induced by the bacterial pathogen *B. thailandensis*. We found that cell-cell fusion occurs within host cell plasma membrane protrusions formed by motile bacteria, with membrane fusion occurring either proximal to the bacterium at the protrusion tip or elsewhere in the protrusion. We also identified TagD5, a component of the T6SS-5 that likely interacts with the T6SS-5 protein VgrG5, as a factor critical for cell-cell fusion. We found that both TagD5 and VgrG5 function at the membrane fusion step of the cell-cell fusion pathway. We further showed that VgrG5 must be secreted within membrane protrusions to support cell-cell fusion. Our results suggest that the T6SS-5 components VgrG5 and TagD5 act within membrane protrusions to promote membrane fusion.

We demonstrated that the first step of the cell-cell fusion pathway is for bacteria undergoing actin-based motility to collide with the host cell plasma membrane and form membrane protrusions that extend from donor cells into recipient cells. Membrane protrusions containing *B. thailandensis* have been observed previously (Kespichayawattana et al., 2000; J. M. Stevens et al., 2005; M. P. Stevens et al., 2005). Our observations further indicate that cell-cell fusion occurs within these protrusions, suggesting that bacterially-induced plasma membrane protrusions function as mediators of cell-cell fusion. Protrusions might mediate key molecular steps leading to cell-cell fusion, such as membrane apposition, membrane fusion, or fusion pore expansion. However, a *B. thailandensis* strain deficient for motility can induce very limited cell-cell fusion (French et al., 2011), indicating that such protrusions, while important, are not absolutely required. Membrane protrusions formed by bacteria undergoing actin-based motility are reminiscent of actin-rich protrusions that promote cell-cell fusion in other contexts, including *Drosophila* myoblast fusion (Kim et al., 2015; Sens et al., 2010), osteoclast fusion (Oikawa et al., 2012), and macrophage fusion (Faust et al., 2019). Force from actin polymerization is also thought to promote virus induced cell-cell fusion by the fusion-associated small transmembrane (FAST) fusogens expressed by a group of nonenveloped, fusogenic reoviruses (Chan et al., 2020, 2021). Similar to these examples, bacterial actin-based motility within protrusions could provide the force necessary to bring neighboring plasma membranes close together, a key step in the cell-cell fusion process (Hernández & Podbilewicz, 2017).

Delineation of the *B. thailandensis*-induced cell-cell fusion pathway at the cellular level also enabled our subsequent analysis of the role of bacterial factors in this process. Because VgrG5 is the only protein known to be secreted by the T6SS-5 and because it is required for cell-cell fusion (Schwarz et al., 2014; Toesca et al., 2014), it is a candidate fusogen protein. Consistent with this idea, we found that VgrG5 must be expressed by a bacterium within a protrusion for cell-cell fusion to occur, placing VgrG5 in the location of the fusion event. Moreover, we found that VgrG5 is required for the membrane fusion step but not for earlier steps in the pathway. Our results are consistent with a direct role for VgrG5 in inducing plasma membrane fusion, although it is possible that VgrG5 does not directly mediate membrane fusion. Ultimately, to define the molecular level mechanism of cell-cell fusion, it will be necessary to demonstrate that the required proteins are sufficient to induce membrane fusion in a minimal system.

We found that an additional component of the T6SS-5, TagD5, is required for membrane fusion. TagD5 (Lennings, West, et al., 2019) is a member of the PAAR family proteins that interact with and are secreted along with VgrG proteins of other T6SS systems (Hachani et al., 2016; Shneider, 2013). This suggests that VgrG5 and TagD5 might form a complex and therefore function together. Consistent with this hypothesis, we showed that TagD5 is required for membrane fusion and that TagD5 contributes to VgrG5 stability or expression. TagD5 is small (119 amino acids) and contains a PAAR structural domain but does not contain a sequence extension present in some PAAR proteins that carries out effector functions (Shneider, 2013; Hachani et al., 2016), so it is unclear how it could directly contribute to membrane fusion. However, VgrG5 contains additional sequences beyond the VgrG structural features that are required for fusion, and therefore it likely carries out effector functions (Pukatzki et al., 2007a; Schwarz et al., 2014; Toesca et al., 2014). A TagD5-VgrG5 complex could act similarly to other PAAR protein-VgrG systems. One particularly relevant example is the *Pseudomonas aeruginosa* Tse6-VgrG1 PAAR protein-VgrG complex which delivers a toxin domain to the cytoplasm of neighboring bacteria during interbacterial competition with only Tse6 contributing effector activity (Quentin et al., 2018; Whitney et al., 2014, 2015). The *P. aeruginosa* Tse6-VgrG1 complex requires chaperones for stability and loading onto the T6SS (Hachani et al., 2016; Quentin et al., 2018; Whitney et al., 2014), suggesting a TagD5-VgrG5 complex may require other yet-to-be-identified bacterial proteins such as chaperones or even secreted effectors. A full understanding of the molecular mechanism of cell-cell fusion will require identification of all the required factors.

Because the T6SS is ejected from bacteria and can puncture neighboring cell membranes (Jurėnas & Journet, 2021), one hypothesis is that this process directly mediates fusion, for example, by disrupting membrane integrity. This hypothesis would predict that membrane fusion occurs proximal to a bacterium. However, we found that membrane fusion does not always occur in close proximity to a bacterium but frequently occurs elsewhere within a membrane protrusion, away from the bacterium. Therefore, our data support a canonical role for the T6SS-5 in secreting effector proteins (Jurėnas & Journet, 2021) rather than in inducing membrane fusion directly. Although our results are insufficient to determine the molecular-level mechanism of membrane fusion during *B. thailandensis* induced cell-cell fusion, they are consistent with membrane fusion involving a canonical fusogen-mediated hemifusion pathway (Hernández & Podbilewicz, 2017). Because we observed that VgrG5 must be expressed within membrane protrusions, we hypothesize that VgrG5 is secreted and released once a bacterium enters a protrusion and then acts within the protrusion to promote cell-cell fusion. This mechanism is consistent with the mechanism of other VgrG proteins that are released upon T6SS secretion (Hachani et al., 2016), including VgrG2b of *P. aeruginosa*, which targets host microtubules (Sana et al., 2015), and VgrG1 of *Vibrio cholerae*, which targets host actin (Ma et al., 2009; Pukatzki et al., 2007b). Therefore, our results are consistent with known functions of the T6SS in secreting bacterial proteins.

In the absence of fusion due to loss of TagD5 or VgrG5, bacteria in protrusions are engulfed by the recipient cell. This pathway is similar to the engulfment of protrusions containing other bacteria, such as *L. monocytogenes* and *R. parkeri*, into double-membrane vacuoles to achieve cell-to-cell spread (Lamason & Welch, 2017). We never observed an engulfed bacterium exiting its double membrane vacuole, and such a defect in accessing the cytosol could explain prior observations that *Bt* Δ*vgrG5* has a growth defect in host cells (Bulterys et al., 2019). Our observations are also consistent with prior observations that protrusion formation through actin-based motility drives bacterial engulfment into the recipient cell (Monack & Theriot, 2001). In addition, we found that *Bt*GFP WT induce cell-cell fusion more quickly than the time it takes for cells to engulf *tagD5*- and *vgrG5*-deficient mutants, indicating that fusion must occur before engulfment occurs. Our work suggests that membrane fusion must be carried out quickly and efficiently to supersede a slower default double membrane protrusion engulfment pathway that is detrimental to the growth and spread of *B. thailandensis*.

Our findings define the cellular level pathway for *B. thailandensis*-induced cell-cell fusion resolving how bacterial motility, bacterial membrane protrusions, and T6SS-5 activity work together to induce cell-cell fusion. Although the T6SS components VgrG5 and TagD5 are directly implicated in membrane fusion, they do not resemble any known fusogens (Podbilewicz, 2014). Therefore, understanding how these proteins function during cell-cell fusion could reveal new insights into membrane fusion mechanisms. The conspicuous length of membrane protrusions formed by *B. thailandensis*, which lend themselves to imaging, makes this a powerful system for continuing to explore the conserved function of membrane protrusions during cell-cell fusion. Continued investigation of this pathway will enhance our understanding of the cellular and molecular mechanisms of membrane fusion and cell-cell fusion.

## Materials and Methods

### Bacterial and mammalian cell culture

*Escherichia coli* strains XL1-blue and BL21(DE3) were obtained from the UC Berkeley MacroLab and were used for plasmid construction and protein expression, respectively. *E. coli* was cultured in liquid or solid lysogeny broth (LB) with or without 100 μg/ml ampicillin or 50 μg/ml kanamycin, when appropriate. *E. coli* RHO3 (López et al., 2009) was grown in LB supplemented with diaminopimelic acid (DAP) (200 mg/ml). *B. thailandensis* E264 was cultured in liquid or solid LB.

Mammalian cell lines (Vero monkey kidney epithelial, RRID:CVCL_0059, HEK293T human embryonic kidney, RRID:CVCL_0045; A549 human lung epithelial, RRID:CVCL_0023; and U2OS human osteosarcoma, RRID:CVCL_0042) were obtained from the University of California, Berkeley Tissue Culture Facility, which authenticated these cell lines prior to freezing, and were not tested for mycoplasma contamination. Cells were grown at 37°C in 5% CO_2_. Vero cells were maintained in DMEM (Invitrogen, 11965-092v) containing 2% fetal bovine serum (FBS, GemCell Bio-Products, 100-500). HEK293T, A549, and U2OS cells were maintained in DMEM containing 10% FBS (Atlas Biologicals, FP-0500-A).

### Plasmid construction

To visualize GFP (Wasabi) in A549 cells, the lentiviral expression vector Wasabi-pIPFCW2 was constructed. The gene encoding Wasabi was amplified by PCR from the plasmid F-tractin-Wasabi-pIPFCW2 (Benanti et al., 2015) with 5’ NheI and 3’ EcoRI cut-sites included in the primer overhangs for subcloning (forward primer 5’ GAACCGTCAGATCCGCTAGCATGGTGAGCAAGGGCG 3’, reverse primer 5’ GGGCGAATTCTTACTTGTACAGCTCGTCCATGC 3’). The F-tractin-Wasabi-pIPFCW2 and amplified *wasabi* gene and were cleaved with NheI and EcoRI and ligated together to produce Wasabi-pIPFCW2.

To make *B. thailandensis* mutants, we used plasmid pEXKm5 (López et al., 2009) for allelic exchange. We PCR-amplified DNA from *B. thailandensis* cells boiled in water. These DNA fragments contained ~500 bp 5’ and 3’ to the region of interest and were subcloned into pEXKm5. To generate the *clpV5-gfp* pEXKm5 plasmid, the 5’ and 3’ends were flanked by sequences in pEXKm5 surrounding the HindIII cut-site. We amplified *clpV5* (primers 5’ CAACGCGCGCAGTAAAGGAGAAGAACTTTTCAC3’and GGGAACTCCTTTATTTGTATAGTTCATCCATGC3’), *gfp* (primers 5’ CAACGCGCGCAGTAAAGGAGAAGAACTTTTCAC3’ and 5’GGGAACTCCTTTATTTGTATAGTTCATCCATGC3’), and ~500 bp 3’ to *clpV5* (primers 5’ TATACAAATAAAGGAGTTCCCGATGTCTTCGTC3’ and 5’CTCGAGGCGGCCGGCTAGCATTGACGATATCGGGAATCG3’). pEXKm5 was digested with HindIII and the 4 fragments were assembled via Gibson cloning (New England Biolabs, E2611S).

To construct the Δ*vgrG5* pEXKm5 plasmid, two fragments were PCR-amplified with 11 bp of homology to each other and this homologous region contained two in-frame stop codons at codon 108 of *vgrG5*. One fragment contained a 5’ XmaI cut-site and ~500 bp 5’ to codon 108 of *vgrG5* (primers 5’ CCCTGTTATCCCTACCCGGGACGCGCGACGCTTCAC 3’ and 5’ GCCTTCCTTCATCAATCGAGATGGCTCTCGTCGTACT 3’) and the other contained ~500 bp 3’ to codon 108 of *vgrG5* and a 3’ XmaI cut-site (5’ TCTCGATTGATGAAGGAAGGCCTCTACTACTACTTCGAGC 3’ and 5’ TCGACTTAAGCCGGCCCGGGGGTGCGCCTGCGAGC 3’). The two fragments were then stitched together via their 11 bp region of homology by overlap PCR.

To construct the Δ*tagD5* pEXKm5 plasmid, two fragments were PCR-amplified. The first fragment contained ~500 bp upstream of the *tagD5* start codon (primers 5’ATCCCTACCCGGGTCGTGCGCATCCGCATCCTCTT 3’ and 5’ GCTCATGCCCGCGCCGCACAGGCCGGAGGCGG 3’) and the second fragment contained ~500 bp downstream of the *tagD5* stop codon (primers 5’GCCTCCGGCCTGTGCGGCGCGGGCATGAGCGATC 3’ and 5’ AGCCGGCCCGGGGATTCGCAGCGGCACGTCGAA 3’). The two fragments were then stitched together via overlap PCR. The stitched fragments and pEXKm5 were digested with XmaI and ligated together.

To express 6xHis-MBP-VgrG5 CTD, we used a version of pETM1 expression vector containing a 6xHis tag, MBP tag, and TEV cleavage site downstream of the SspI cut site. A fragment of *vgrG5* encoding a C-terminal domain of *vgrg5* (aa718-1012, *vgrG5-ctd*) was amplified by PCR from *B. thailandensis* (primers 5’ ACCTGTACTTCCAATCCAATCGCACGCTGCTCTCGAAAATC 3’ and 5’ ATCCGTTATCCACTTCCAATGCCTAGCTGGATCAACTGTC 3’) and subcloned into the SspI site of the pETM1.

To express 6xHis-SUMO-VgrG5-CTD, *vgrG5-ctd* was amplified by PCR (primers 5’ACCTGTACTTCCAATCCAATCGCACGCTGCTCTCGAAAATC 3’ and 5’ ATTGGAAGTGGATAACGGATGCCTAGCTGGATCAACTGTC 3’) and subcloned into the SspI site of plasmid pSMTp3, 3’ to the portion encoding HIS-SUMO.

### *B. thailandensis* strain construction

*B. thailandensis* strains were created by allelic exchange, as previously described (Benanti et al., 2015; López et al., 2009). Bi-parental matings between *B. thailandensis* strain E264 and *E. coli* RHO3 (López et al., 2009) harboring a pEXKm5 derivative were performed to introduce pEXkm5 into *B. thailandensis*, followed by selection on 50 μg/ml kanamycin-containing plates that lacked DAP to select against *E. coli* RHO3. Uptake of pEXKm5 was also confirmed by PCR detection of the *sacB* gene. The integrated vector backbone was removed by growth in non-selective YT media (5 g/l yeast extract (VWR, EM1.03753.0500), 5 g/l tryptone (Fisher Scientific, BP1421-500)) and screening for loss of b-glucuronidase activity via plating on YT plates containing 50 μg/ml X-Gluc (cyclohexlammonium salt, Gold Biotechnologies, G1281C1). Strains were confirmed by PCR amplification and DNA sequencing of the region of interest.

### Transient transfections, transduction, and cell line production

For retroviral transduction to visualize GFP (Wasabi) in A549 cells, viral particles were packaged by transfecting HEK293s plated 24 h prior at 5×10^5^ cells/well (2 ml/well, 6-well plate), via calcium phosphate transfection with 750 ng pMDL-RRE, 450 ng pCMV-VSVg, 300 ng RSV-Rev and 1500 ng Wasabi-pIPFCW2. Approximately 22 h after transfection, the media was replaced with 2 ml fresh media. After an additional 21 h, the supernatant, which contains viral particles, was collected from each well, and cell debris was cleared by filtration through a 0.45 μm syringe filter. The viral supernatant was added to A549 cells and polybrene (Santa Cruz Biosciences, sc-134220) was added to 10 μg/ml to enhance the infection efficiency. After transduction, fresh media was added at 24 h post infection (hpi), and at 48 hpi cells transduced with Wasabi-pIPFCW2 were selected with 3-4 mg/ml puromycin (Calbiochem, 540411) and sorted for mid-range expression of Wasabi.

### Bacterial Infections of host cells

*B. thailandensis* strains (Δ*motA2;ClpV5-GFP* (*Bt*GFP WT) (this study), Δ*motA2*; Δ*vgrG5* (Δ*vgrG5*) (this study), *Bt*GFP Δ*vgrG5* (this study), Δ*motA2*; Δ*tagD5* (Δ*tagD5*) (this study), *Bt*GFP Δ*tagD5* (this study), Δ*motA2;BFP* (*Bt*BFP) (Benanti et al., 2015)) were streaked from frozen stocks onto LB agar plates. Bacteria were swabbed from plates to inoculate LB liquid media and were grown with shaking at 37°C for 3-16 h. Prior to infections, the OD_600_ of cultures was measured in order to calculate the number of bacteria to infect with (OD_600_ of 1 =5×10^8^ cfu/ml) to achieve the proper multiplicity of infection (MOI). Bacterial cultures were pelleted and resuspended in PBS (ThermoFisher, 10010049. Composition: Potassium Phosphate monobasic (KH2PO4), 1.0588236mM; Sodium Chloride (NaCl), 155.17241mM; Sodium Phosphate dibasic (Na2HPO4-7H2O), 2.966418mM). Mammalian cells were seeded at least 24 h before infection and immediately prior to infection were washed with PBS and provided with fresh DMEM with 10% FBS. Bacteria were added directly to media on cells, media was pipetted or rocked gently to mix, and bacteria were left to invade for 45 min to 1 h at 37°C unless otherwise stated. Cells were rinsed once with PBS, then DMEM with 10% FBS and 0.5 mg/ml gentamicin (Fisher Scientific, MT30-005-cR) was added. For mixed-strain infections, infections lasted longer, as detailed below.

For live cell imaging of spread, confluent monolayers of A549 cells were infected. For infection with *Bt*GFP WT (Δ*motA2;clpV5-GFP*), a mix of A549 TagRFP-T-farnesyl (Lamason et al., 2016) and A549 GFP cells at a 1:1 ratio (6×10^5^ cells/dish) were plated in 20 mm MatTek dishes (Mat Tek Corp., P35G-1.5-20-C). Cells were infected as described above at an MOI of 10-50 and imaged at 12-18 h. For live imaging of *Bt*GFP Δ*vgrG5* and *Bt*GFP Δ*tagD5*, A549 TagRFP-T-farnesyl were plated in 20 mm Mat Tek dishes (6×10^5^ cells/dish) 24-48 h before infection. Cells were infected as described above at an MOI of 100 and imaged at 24-30 hpi.

For co-infections of *Bt*BFP WT and *Bt*GFP Δ*vgrG5*, a mix of A549 TagRFP-T-farnesyl and A549 GFP cells at a 1:1 ratio or 4:1 were plated in 20 mm Mat Tek dishes (6×10^5^ cells/dish) at least 24 h before infection. Infections were done two ways. For two videos, *Bt*BFP WT were infected first at an MOI of 100, and allowed to invade for 2 h. Then at 5 h after the initial infection, *Bt*GFP Δ*vgrG5* were added at an MOI of 100 and allowed to invade for 2 h. Imaging was performed at 9-12 hpi. For the other eight videos, *Bt*BFP WT and *Bt*GFP Δ*vgrG5* were infected simultaneously, allowed to infect for 1.5 h, and imaging was performed at 14-19 hpi. For some of the experiments, each strain was used to infect at an MOI of 50 and in others, they were used to infect at an MOI of 20 (*Bt*BFP WT) and 80 (*Bt*GFP Δ*vgrG5*).

### Live cell imaging

Before imaging, infected cells in 20 mm Mat Tek dishes were washed once with PBS before addition of 1.5 ml FluoroBrite DMEM Media (Invitrogen, A18967-01) supplemented with 10% FBS and 1XGlutaMAX (Gibco, 35050-061) and 0.5 mg/ml gentamycin.

Images were captured on a Nikon Ti Eclipse microscope with a Yokogawa CSU-XI spinning disc confocal, 60X (1.4 NA) Plan Apo objective, a Clara Interline CCD Camera (Andor Technology), and MetaMorph software (Molecular Devices). 3 image Z-stacks were captured at 15 s intervals for 30-90 m. For mixed infections, images were taken every 20 s. Images were processed using ImageJ (Version 2.1.0/1.53c) and assembled in Adobe Illustrator (version 25.3.1). Spread events were then observed and a dataset was collected of individual spread events in which we were able to identify which bacterium induced cell-cell fusion. The kinetics and membrane morphology for each spread event were recorded. Maximum protrusion was defined as the longest protrusion length observed before earliest sign of GFP diffusion into the recipient cell (for *Bt*GFP WT) or engulfment (for *Bt*GFP Δ*vgrG5* and *Bt*GFP Δ*tagD5*). Time of spread was defined as the time of protrusion entry to the earliest sign of GFP diffusion into the recipient cell (for *Bt*GFP WT) or engulfment (for *Bt*GFP Δ*vgrG5* and *Bt*GFP Δ*tagD5*).

### Plaque assay

For plaque assays, Vero cells were plated in 6-well plates (6×10^5^ cells/well), infected at an MOI of 2, and bacteria were allowed to invade for 45 min. Infected cell monolayers were washed once with PBS and overlayed with 3 ml of 0.7% agarose in DMEM with 5% FBS and 0.5 mg/ml gentamycin. At 31 hpi, 1 ml of 0.7% agarose in PBS containing neutral red (Sigma, N6264) at 1:20 dilution was overlayed onto wells (final concentration on cells was 1%). 14 h after addition of neutral red, plates were scanned and plaque area was measured using ImageJ (Version 2.1.0/1.53c).

### Protein expression and purification

For expression of VgrG5 in broth culture (J. Wong et al., 2015), *B. thailandensis* strains were grown overnight and then diluted 1:10 in 3.5 ml LB. After 2 h, cultures were split into 2 tubes with 1.5 ml each and L-Glutathione reduced (GSH, Sigma-Aldrich, G4251) was added to 50 mM in one of them. Cultures were grown for 2 h followed by processing for western blotting as described below.

To generate the anti-VgrG5 antibody, 6xHis-MBP-TEV-VgrG5-CTD was expressed in *E. coli* BL21. Protein expression was induced with 1mM IPTG at 37°C 1 h. Cells were pelleted at 4539.5 xg and resuspended in 50 mM Tris HCl pH 8.0, 200 mM KCl, 1 mM EDTA, and protease inhibitors (1 μg/ml each leupeptin (MilliporeSigma, L2884), pepstatin (MilliporeSigma, P5318), chymostatin (MilliporeSigma, E16), 1 mM phenylmethylsulfonyl fluoride (PMSF, 600 MilliporeSigma, 52332)) and stored at −80°C. Cells were thawed, imidazole was added to 5 mM, and cells were incubated with 1 mg/ml lysozyme (Sigma, L4919-5G) for 15 min on ice and then sonicated at 4°C (6× 12 s pulses, 50% power). The lysate was spun at 20198 xg, 4°C, for 25 min. The supernatant was incubated for ~2 h rotating at 4°C with Ni-NTA Resin (Qiagen, 1018244) that had been washed with wash buffer (20mM Tris HCl pH 8.0, 200 mM NaCl, 1mM DTT, 20mM imidazole). Cleared lysate was incubated with resin for ~2 h, rotating at 4°C, and resin was washed with 3 ml wash buffer. Protein was eluted stepwise in 50 mM, 200 mM, and 500 mM imidazole. Elutions containing VgrG5-CTD were desalted using Amicon Ultra-4 Centrifugal Filter Units (Merck Millipore Ltd., UFC801096) into 10 mM imidazole, 50 mM Tris HCl pH 8.0, 200 mM NaCl, and incubated overnight at 4°C with TEV protease at a VgrG5-CTD:TEV ratio of 1:100. MBP-VgrG5-CTD was run over an Ni-NTA column as described above but with 10 ml wash buffer containing 30 mM imidazole. The wash was collected in 1 ml fractions. The rest of the protein was eluted in elution buffer containing 200 mM imidazole. The washes and elution were pooled and then concentrated to 1 mg using a desalting column. This resulted in a mixed population of mostly uncleaved 6xHis-MBP-VgrG5-CTD and some VgrG5-CTD.

For antibody affinity purification, HIS-SUMO-VgrG5-CTD was expressed in bacteria as described above and purified using Ni NTA resin as described above but eluted with 200 mM imidazole. The protein was then further purified by concentrating and running over a gel filtration column (CYZ superdex 200 increase, Sigma) in 20 mM HEPES pH 8.0, 200 mM NaCl. Fractions containing HIS-SUMO-VgrG5-CTD were pooled and concentrated as described above to 1 mg/ml.

### Antibody production, purification, and validation

To generate rabbit-anti VgrG5 antibodies, purified VgrG5-CTD protein was used to inoculate rabbits at Pocono Rabbit Farm and Laboratory (Canadensis, PA) where a 91-day custom antibody protocol was performed.

To purify the anti-VgrG5 antibody, purified HIS-SUMO-VgrG5-CTD was concentrated to 0.5 ml and was combined with 0.5 ml coupling buffer (200 mM NaHCO3 pH 8.3, 500 mM NaCl). This was then coupled onto NHS-ester Sepharose 4 Fast Flow resin (GE Healthcare, 17-0906-01) for 4 h at 4°C. 10 ml of serum was diluted 1:1 in binding buffer (20 mM Tris, pH 7.5), 0.2 μm filtered, and rotated for 1 h at room temperature with the resin. After washing with binding buffer, the antibody was eluted off of the resin with 100 mM glycine, pH 2.5, and 1 ml fractions were collected. Eluted fractions were immediately neutralized with 1 M Tris pH, 8.8 to 65.4 mM final concentration. Elutions that recognized VgrG5 via western blot (elutions 2 and 3) were pooled and dialyzed into 20 mM HEPES, pH 7.0, 100mM NaCl. Aliquots were flash-frozen and stored at −80°C or supplemented with 35% glycerol and stored at −20°C. To validate the VgrG5 antibody, western blots were performed as described below.

### Western blotting

For detection of VgrG5 in glutathione-induced samples, 100 μl of broth culture was washed once with PBS and boiled in 1X SDS sample buffer three times for 10 min each. Samples were run on a 10% SDS-PAGE gel and the gel contents were transferred to a nitrocellulose membrane (ThermoFisher, 88018). Membrane was blocked for 30 min in TBS-T (20 mM Tris pH 8.0, 150 mM NaCl, 0.1% Tween 20 (Sigma, P9416) containing 5% milk (Genesee, 20-241), then incubated with 1:5000 anti-VgrG in 5% milk in TBS-T overnight at 4°C. The membrane was then washed 3 × 5 min in TBS-T and incubated with 1:5000 goat anti-rabbit HRP secondary antibody (Santa Cruz Biotechnology, sc-2004) in 5% milk in TBS-T for 1 h followed by 3 × 5 min washes in TBS-T. To detect secondary antibodies, ECL HRP substrate kit (Advansta, K-12045) was added to the membrane for 1 min at room temperature and developed using HyBlot ES High Sensitivity Film (Thomas Scientific 1156P37).

### Statistics and sample size

Statistical analyses were performed using GraphPad PRISM v.9. Statistical parameters and significance are reported in the Figure Legends. Comparisons were made using unpaired, two-tailed Mann-Whitney tests and differences were considered to be statistically significant when P < 0.05, as determined by an unpaired Mann-Whitney test. For comparing time to engulfment across different strains, a one-way ANOVA test with multiple comparisons was performed. Sample size of n = 20 independent events was selected for cell-cell fusion events and cell-cell engulfment spread events based on the experimental limitations of capturing such rare events. Protrusion lengths could not be measured during all cell-cell fusion events, resulting in a smaller sample size for those datasets. Engulfment of *Bt*GFP WT events were extremely rare, resulting in a smaller number of events observed and a smaller dataset. For live cell imaging experiments, each imaging session was a biological replicate without technical replicates. The sample size for plaque size assays was determined by the number of plaques present in two wells of a 6 well dish (9-11 plaques for *Bt* WT, 0 for *Bt* Δ*vgrG5* and *Bt* Δ*tagD5*). Each plaque measured was a technical replicate with n = 3 biological replicates performed. There was no randomization or blinding.

## Abbreviations

T6SS: type VI secretion system
MNGC: multinucleated giant cell
PAAR: proline-alanine-alanine-arginine
*Bt*: *Burkholderia thialandensis*

## Acknowledgements

We thank previous Welch Lab members whose work supported this project, including Erin Benanti, Joseph Graham, Curtis Sera, Rebecca Lamason, Jessica Leslie, and Caroline Boyd, as well as current lab members, for critical feedback throughout the development of this project. We also thank the following UC Berkeley core facilities and their facility members for providing equipment, reagents, and technical support to complete this work: Holly Aaron and Feather Ives (CRL Molecular Imaging Center); and Alison Killilea (Cell Culture Facility). We thank David Drubin, Sarah Stanley, and Daniel Fletcher for technical discussion and critical guidance for this work. We also thank Neil Fischer for proofreading the manuscript. This work was funded by grant R35 GM127108 from the NIH/NIGMS to M.D.W.

---

**Supplemental Figure 1:**
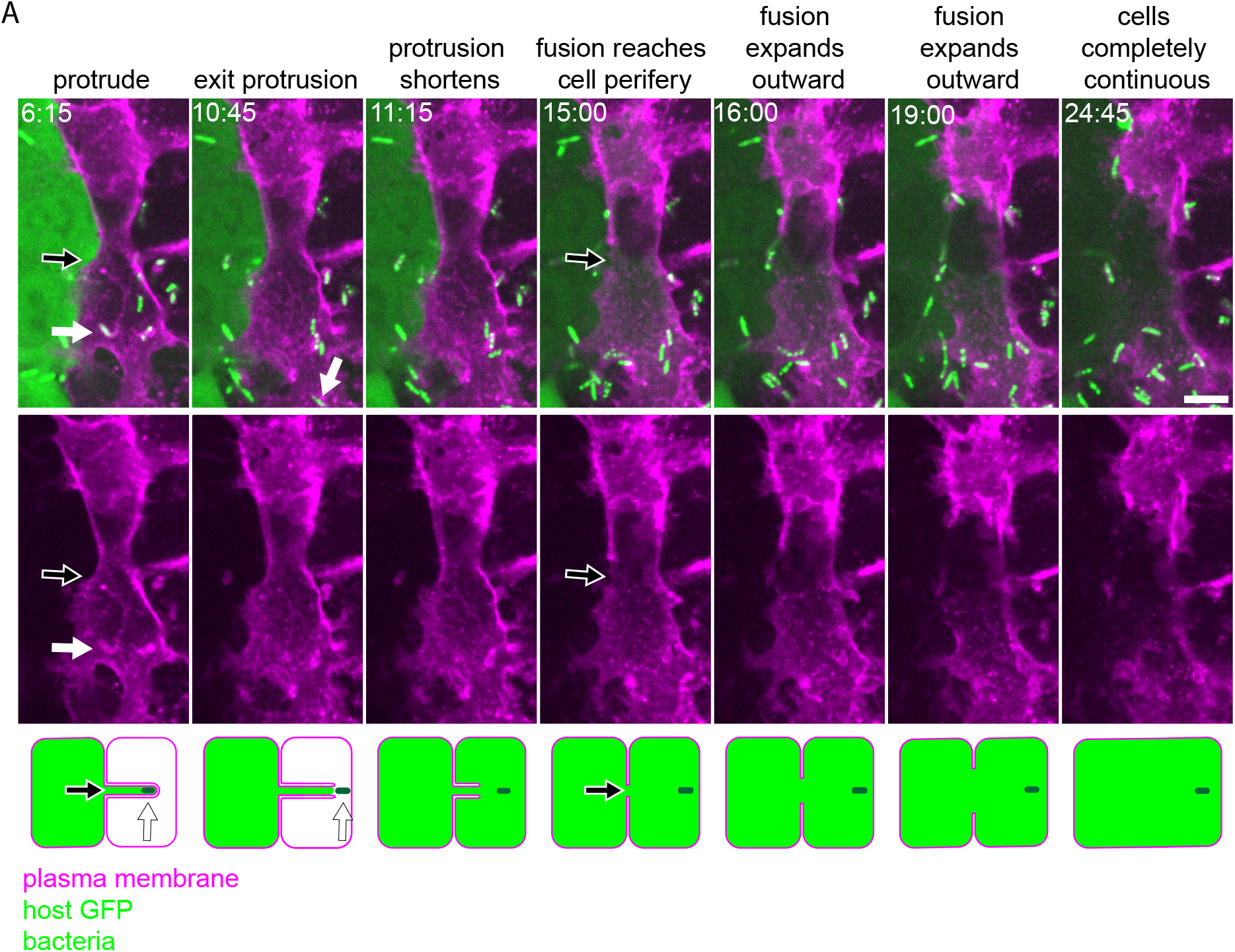
Details of *B. thailandensis* inducing cell-cell fusion expanding from a membrane protrusion. (A-C) Live-cell imaging stills of two examples of *Bt*GFP WT while inducing cell-cell fusion. A 1:1 mixture of A549 cells that expressed the plasma membrane marker TagRFP-T-farnesyl or cytoplasmic GFP were used. Times represent min:s post protrusion formation. Images taken at ~16 h post infection. Scale bars are 5 μm. White arrows highlight the bacterium forming the protrusion. Black arrows highlight the region of protrusion entry. Note that this movie is not part of the quantification dataset because the protrusion formed before the movie began.

**Supplemental Figure 2:**
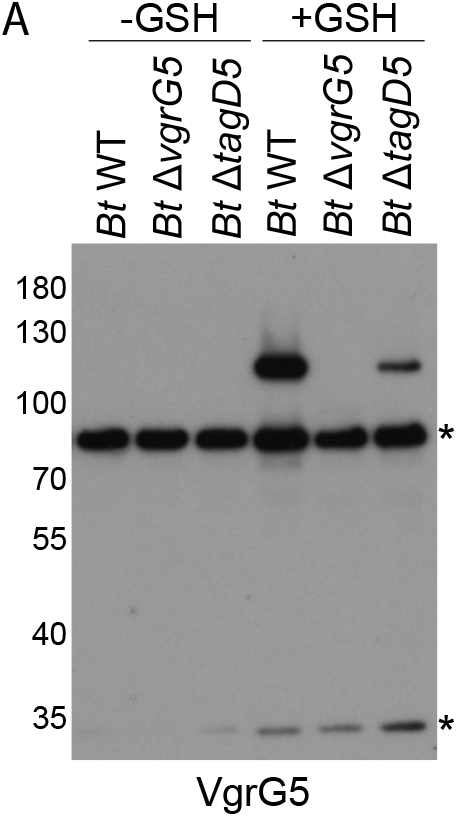
VgrG5 expression in glutathione-induced *Bt* WT, *Bt* Δ*vgrG5* and *Bt* Δ*tagD5*. (A) Western blot of cell lysates from the indicated strains grown in liquid cultures induced or uninduced with glutathione (GSH). Blot was probed with anti-VgrG5 antibodies. The asterisks indicate non-specific bands.

**Supplemental Figure 3:**
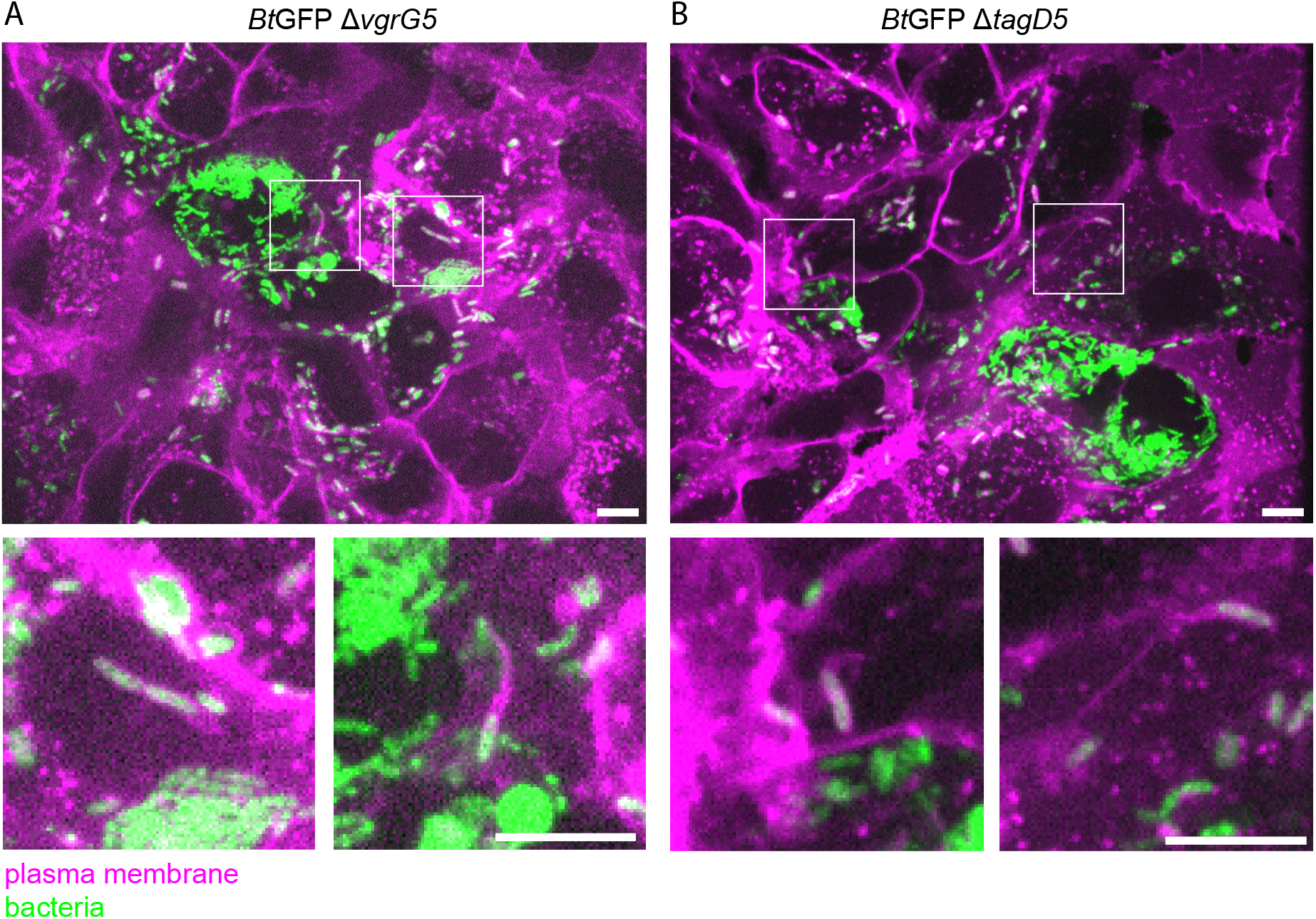
*Bt*GFP Δ*vgrG5* and *Bt*GFP Δ*tagD5* can form protrusions from secondary cells. (A and B) Live imaging stills showing protrusions formed by motile bacteria in secondary cells after spread. Host cells are A549 cells expressing TagRFP-T-farnesyl. Images taken at ~24 h post infection with *Bt*GFP Δ*vgrG5* or *Bt*GFP Δ*tagD5*. Scale bars are 5 μm.

**Supplemental Figure 4:**
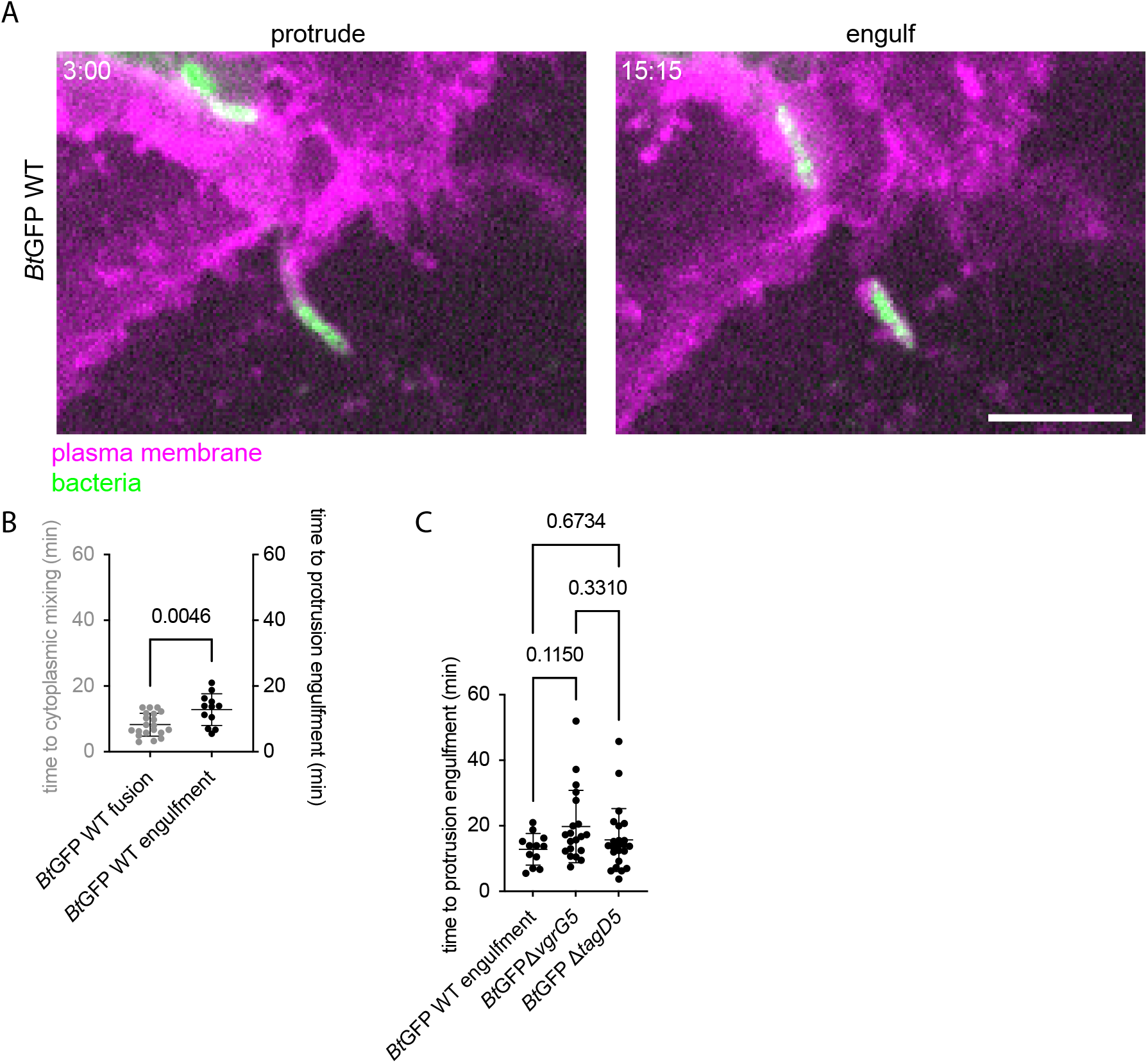
Quantification of engulfment of *Bt*GFP WT, *Bt*GFP Δ*vgrG5* and *Bt*GFP Δ*tagD5*. (A) Live cell imaging stills of *Bt*GFP WT during engulfment into recipient cells. A549 cells that expressed TagRFP-T-farnesyl were used. Times represent min:s post protrusion formation. Images taken at ~16 h post infection. Scale bar is 5 μm. (B) Time to cytoplasmic mixing (n = 20) or protrusion engulfment for *Bt*GFP WT (n = 14). P value was calculated by an unpaired Mann-Whitney test. (C) Time to engulfment of *Bt*GFP WT (n = 14), *Bt*GFP Δ*vgrG5* (n = 20), or *Bt*GFP Δ*tagD5* (n=23). P = 0.1172, calculated by one-way ANOVA with Tukey’s multiple comparisons test. Data are mean +/− SD.

## Video Legends

**Video S1. An example of *B. thailandensis* inducing cell-cell fusion at the tip of plasma membrane protrusions.** Confocal timelapse imaging of *Bt*GFP WT inducing cell-cell fusion following infection of a 1:1 mixture of A549 cells that expressed either the plasma membrane marker TagRFP-T-farnesyl or cytoplasmic GFP. Cells were imaged every 15 s starting at ~12-18 h post infection. Video is shown at 5 frames per s. Scale bar is 5 μm. Time 0:00 marks the moment the bacterium began forming a protrusion. This video accompanies Figure 1A.

**Video S2. An example of *B. thailandensis* inducing cell-cell fusion at the tip of plasma membrane protrusions.** Confocal timelapse imaging of *Bt*GFP WT inducing cell-cell fusion following infection of a 1:1 mixture of A549 cells that expressed either the plasma membrane marker TagRFP-T-farnesyl or cytoplasmic GFP. Cells were imaged every 15 s starting at ~12-18 h post infection. Video is shown at 5 frames per s. Scale bar is 5 μm. Time 0:00 marks the moment the bacterium began forming a protrusion. This video accompanies Figure 1A.

**Video S3. An example of *B. thailandensis* inducing cell-cell fusion within the protrusion but not at its tip.** Confocal timelapse imaging of *Bt*GFP WT inducing cell-cell fusion following infection of a 1:1 mixture of A549 cells that expressed the plasma membrane marker TagRFP-T-farnesyl or cytoplasmic GFP. Cells were imaged every 15 s starting at ~12-18 h post infection. Video is shown at 5 frames per s. Scale bar is 5 μm. Time 0:00 marks the moment the bacterium began forming a protrusion. This video accompanies Figure 1C.

**Video S4. An example of *B. thailandensis* inducing cell-cell fusion within the protrusion but not at its tip.** Confocal timelapse imaging of *Bt*GFP WT inducing cell-cell fusion following infection of a 1:1 mixture of A549 cells that expressed the plasma membrane marker TagRFP-T-farnesyl or cytoplasmic GFP. Cells were imaged every 15 s starting at ~12-18 h post infection. Video is shown at 5 frames per s. Scale bar is 5 μm. Time 0:00 marks the moment the bacterium began forming a protrusion. This video accompanies Figure 1C.

**Video S5. VgrG5 acts at the membrane fusion step.** Confocal timelapse imaging of *Bt*GFP Δ*vgrG5* during cell-to-cell spread following infection of A549 cells that expressed the plasma membrane marker TagRFP-T-farnesyl. Cells were imaged every 15 s starting at ~24 h post infection. Video is shown at 5 frames per s. Scale bar is 5 μm. Time 0:00 marks the moment the bacterium began forming a protrusion. This video accompanies Figure 2A.

**Video S6. Engulfment of *Bt*GFP WT.** Confocal timelapse imaging of *Bt*GFP WT in a 1:1 mixture of A549 cells that expressed the plasma membrane marker TagRFP-T-farnesyl or cytoplasmic GFP. Cells were imaged every 15 s starting at ~14-19 h post infection. Video is shown at 5 frames per s. Scale bar is 5 μm. Time 0:00 marks the moment the bacterium began forming a protrusion. This video accompanies Supplemental Figure 3A.

**Video S7. *B. thailandensis* must secrete VgrG5 within a protrusion to induce cell-cell fusion.** Confocal timelapse imaging of *Bt*GFP Δ*vgrg5* spreading from an MNGC initially formed by cell-cell fusion induced by *Bt*BFP WT bacteria that expressed both TagRFP-T-farnesyl and cytoplasmic GFP. *Bt*GFP Δ*vgrg5* is forming a protrusion that extends into an unfused cell that only expressed TagRFP-T-farnesyl. Cells were imaged every 20 s starting at ~24 h post infection. Video is shown at 5 frames per s. Scale bar is 5 μm. Time 0:00 marks the start of image acquisition. This video accompanies Figure 3B.

**Video S8. TagD5 acts at the membrane fusion step.** Confocal timelapse imaging of *Bt*GFP Δ*tagD5* during cell-to-cell spread in A549 cells that expressed the plasma membrane marker TagRFP-T-farnesyl. Cells were imaged every 15 s starting at ~24 h post infection. Video is shown at 5 frames per s. Scale bar is 5 μm. Time 0:00 marks the moment the bacterium began forming a protrusion. This video accompanies Figure 4B.

